# Adolescent Development of Biological Rhythms: Estradiol Dependence and Effects of Combined Contraceptives

**DOI:** 10.1101/2021.07.20.453145

**Authors:** Azure D. Grant, Linda Wilbrecht, Lance J. Kriegsfeld

**Author notes:** Address Correspondence to: Lance J. Kriegsfeld, PhD, Department of Psychology, Integrative Biology, Graduate Group in Endocrinology and The Helen Wills Neuroscience Institute, 2121 Berkeley Way West, University of California, Berkeley, CA, 94720, United States.

## Abstract

**Purpose:** Adolescence is a period of continuous development, including the maturation of endogenous rhythms across systems and timescales. Although these dynamic changes are well recognized, their continuous structure and hormonal dependence have not been systematically characterized. Given the well-established link between core body temperature (CBT) and reproductive hormones in adults, we hypothesized that high-resolution CBT can be applied to passively monitor pubertal development and disruption with high fidelity.

**Methods:** To examine this possibility, we used signal processing to investigate the trajectory of CBT rhythms at the within-day (ultradian), daily (circadian), and ovulatory timescales, their dependence on estradiol, and the effects of hormonal contraceptives.

**Results:** Puberty onset was marked by a rise in fecal estradiol (fE2), followed by an elevation in CBT and circadian power. This time period marked the commencement of 4-day rhythmicity in fE2, CBT, and ultradian power marking the onset of the estrous cycle. The rise in circadian amplitude was accelerated by E2 treatment, indicating a role for this hormone in rhythmic development. Contraceptive administration in later adolescence reduced CBT and circadian power and resulted in disruption to 4-day cycles that persisted after discontinuation.

**Conclusions:** Our data reveal with precise temporal resolution how biological rhythms change across adolescence and demonstrate a role for E2 in the emergence and preservation of multiscale rhythmicity. These findings also demonstrate how hormones delivered exogenously in a non-rhythmic pattern can disrupt rhythmic development. These data lay the groundwork for a future in which temperature metrics provide an inexpensive, convenient method for monitoring pubertal maturation and support the development of hormone therapies that better mimic and support human chronobiology.

## Introduction

Adolescence is a period of rhythmic reorganization during which physiology transitions from a non-reproductive juvenile state into reproductive early adulthood(1–5). Historically, researchers have relied on self-report, tanner staging, or infrequent salivary or blood hormone samples to track milestones of pubertal development. We reasoned that a time series characterization of continuous core body temperature (CBT)(6–10) could reflect dynamic, rhythmic features of hormonal development across the pubertal trajectory. If feasible, CBT could become a highly convenient and more accurate pubertal staging tool with unprecedented temporal resolution. Its application could also improve our understanding of the range of typical pubertal trajectories and factors that drive deviation from these trajectories.

Rhythmic development occurs at multiple timescales, including within-a-day (ultradian rhythms; URs)(11), daily (circadian rhythms; CRs)(12, 13), and multi-day ovulatory cycles in females (ovulatory rhythms; ORs)(14). These rhythms occur across physiological systems, serving to increase the efficiency of signal transduction(15–17), temporally segregate incompatible processes(18), synchronize internal systems to the environment(19), and maximize reproductive success(20). Although structure at one timescale can be modulated by changes at another(21, 22), distinct mechanisms underlie each rhythmic frequency. This suggests individual rhythmic frequencies may each follow distinct developmental trajectories.

Coordinated URs are observed in hypothalamic-pituitary-peripheral axes and beyond in adult mammals, with broad manifestation in systems including cardiovascular outputs, thermoregulation, and even cognition (reviewed in(6,23–26)). Many URs, including those in reproductive and growth hormones, are present early in life(27) and increase in amplitude from pre-to-mid puberty(28–30). Some URs increase markedly in frequency and amplitude around puberty onset (e.g., gonadotropin releasing hormone (GnRH) luteinizing hormone (LH) and estradiol (E2)(11,28,31–35),(28, 35)). In adults, URs are modulated by time of day(36) and phase of the ovulatory cycle(21), suggesting that pubertal modifications at the CR and OR timescales likely impact UR structure. Although URs are thought to be centrally controlled, potentially via interaction of dopaminergic and hypothalamic circuits(23,37–42), relatively little is understood about mechanisms and progression of URs across adolescence.

Whereas the mechanisms and phenomenology of ultradian development require much further study, those underlying circadian rhythms and pubertal changes to CRs are well documented. Circadian rhythms are nearly ubiquitous, generated intracellularly via interlocked transcription-translation feedback loops, and are governed by a central pacemaker within the suprachiasmatic nucleus (SCN) of the hypothalamus(43–45). Although changes in SCN output and connectivity during adolescence are not well characterized, integration of new neurons into the central clock(46) and reproductive neurocircuitry(47) alongside adolescent increases in SCN input to the GnRH system(48) may contribute to downstream CR changes. For example, circadian amplitude appears to increase across puberty in many systems (e.g., cortisol(49), activity(45), and potentially temperature(50)), while emerging for the first time in others (e.g., LH(34) and FSH(29)). Finally, circadian activity(45) and sleep-wake(51) rhythms are phase delayed during puberty(52) and are more vulnerable to disruption by mistimed light and food cues than in adults(53).

In contrast to ultradian and circadian rhythms, which are apparent to variable degrees in juveniles, the female ovulatory, or estrous, cycle emerges for the first time in adolescence(14). Briefly, in spontaneously-ovulating rodents, the 4 to 5 day cycle begins with rising E2 levels that maintain LH at low concentrations through negative feedback. When E2 is sufficiently high, and a pool of ovarian follicles has matured, E2 positive feedback integrates with circadian signaling and progesterone of neural origin(47,54,55) to induce a preovulatory LH surge that initiates ovulation(55–60). Subsequent formation of the corpus luteum leads to a brief rise in circulating progesterone prior to beginning the next cycle. As with URs and CRs, ORs manifest as changes in a number of systems, including metabolic hormones, cardiac output, and thermoregulation(61–63). Although it is clear that ORs emerge at puberty, the continuous patterns of commencement and stabilization are poorly understood. Pre-pubertal ovarian follicles typically undergo development and atresia without substantial release of sex steroids(30). Soon after the emergence of the first cycle, at menarche in girls(20), cycles have a higher likelihood of anovulation or low post-ovulatory progesterone compared to adults(64). Although high temporal resolution patterns are unknown, large increases in plasma E2 and FSH occur from pre to mid puberty(28,34,65). Given these changes in hormones across puberty, one aim of the present investigation was to employ continuous, longitudinal, and high-resolution CBT monitoring alongside daily E2 measures to characterize the emergence of the ovulatory cycle in rats.

By characterizing rhythmic outputs that reflect underlying physiological change across adolescence, a greater understanding of typical progression can be garnered and the impact of exogenous hormone manipulation on temporal trajectories can be observed. Temporal disruption at all three timescales is associated with negative health outcomes in adults(66–71), and adolescence may be a sensitive period where disruptions have rapid(1) and potentially long-term health impacts(2,53,72,73). A growing proportion of teenage girls (estimated between 22-54% across the first two decades of the 21^st^ century(74, 75)) receive hormonal contraceptives for a variety of purposes, including pregnancy prevention(76) or treatment of menstrual symptoms(77), and acne(78). These hormonal contraceptives aim to prevent ovulation, sperm penetration, and implantation by maintaining elevated progestin levels akin to the post-ovulatory portion of the cycle (with or without estrogen derivatives)(79–83). As they are delivered at static or once daily bolus concentrations that differ from the endogenous, multiscale rhythmic pattern of release(84–86), hormonal contraceptives can be considered a form of temporal endocrine disruption(87, 88). Hormonal contraceptive use is associated with elevated body temperature(89), decoupling of follicular maturation cycles within the ovary(88, 90), weight change(91, 92), mental health risks(93–95), lasting luteal phase deficiency(96), and a variety of other off target effects(97–100). Furthermore, women under 21 are more likely to exhibit anovulatory cycles following birth control cessation than are older indivdiuals(101), suggesting that contraceptives taken during late adolescence may be more disruptive than in adulthood. Although currently considered safe, discontinuation rate is high(102) and impact on the temporal progression of development is unclear.

CBT as a continuous surrogate marker for pubertal status may provide convenience and new information to longitudinal research studies on this topic and beyond. Continuous measures of activity have been used for rhythmic monitoring(52), but this measure is not as closely tied to high-frequency changes driven by hormonal systems(9, 10) and does not appear to predict individual health events with comparable specificity(9,10,103). In contrast, CBT exhibits clear URs, CRs, and ORs that reflect underlying hormonal changes(6,61,104), including ultradian rhythmic CBT patterns that mirror LH(21, 105) and estradiol(21) prior to ovulation. Likewise, CBT is a reliable phase marker for circadian rhythmicity(106, 107). Hormonal alterations across the ovulatory cycle directly influence CBT, with E2 decreasing temperatures prior to ovulation, and E2 with progesterone increasing temperature following ovulation(8, 106). As a result, we hypothesized that insight into changes in the temporal structure of underlying physiology across adolescence can be garnered via assessment of continuous CBT levels and rhythms.

The present study employed continuous CBT to characterize rhythmic change across adolescent development and examine the role of pubertal onset of E2 production in guiding the typical developmental trajectory. Additionally, because late pubertal contraceptive use might act to disrupt the typical progression of rhythmic developmental changes, we examined the impact of a common contraceptive regimen (i.e., ethinyl estradiol and levonorgestrel) on endogenous estradiol concentrations and CBT rhythms. As the pulse amplitude of multiple hormones increases across adolescence, we hypothesized that the amplitude of CBT URs would be similarly impacted. We also hypothesized that CR amplitude and overall body temperature would increase during adolescence, and that these increases would be influenced by E2. As changes to rhythmicity have primarily been reported from early to mid adolescence, we hypothesized that rhythmic restructuring would be most pronounced during this period. Finally, we hypothesized that rhythmic patterns of body temperature change identified during adolescent development would be disrupted during and potentially after, the cessation of contraceptive administration.

## Materials and Methods

### Animals

Female Wistar rats were purchased at 250 g from Charles River (Charles River, Wilmington, MA) and bred in the lab and weaned at p21. Weanlings were housed in standard translucent propylene (96 x 54 x 40 cm) rodent cages, and provided *ad libitum* access to food and water, wood chips for floor cover, bedding material, and chew toys for the duration of the study. To minimize social isolation stress, which is known to affect pubertal development(108, 109), animals were housed with a same sex, non-experimental sibling. Animals were maintained on a 12:12 light dark (LD) cycle; light intensity during the photo- and scotophases were ∼500 lux white light and <1 lux red light, respectively, with lights on at 1 AM and off at 1 PM. Animals were gently handled before weighing daily to minimize stress. To prevent mixing of feces used for hormone analysis, cage mates were separated by a flexible stainless steel lattice that permitted aural, scent, and touch interaction between siblings. A total of 64 animals were included in the study: 32 as experimental animals (Intact, Intact + Contraceptives, Ovariectomized (OVX) and OVX + E2; n=8/group), and 32 as social, littermate partners. All procedures were approved by the Institutional Animal Care and Use Committee of the University of California, Berkeley.

### Core Body Temperature Data Collection

Data were gathered with G2 E-Mitter implants that chronically record CBT (Starr Life Sciences Co., Oakmont, PA). At weaning, G2 E-Mitters were implanted in the intraperitoneal cavity under isoflurane anesthesia with analgesia achieved by subcutaneous injections of 0.03 mg/kg buprenorphine (Hospira, Lake Forest, IL) in saline (administered every 12 h for 2 days after surgery). E-Mitters were sutured to the ventral muscle wall to maintain consistent core temperature measurements. Recordings began immediately, but data collected for the first 4 days post-surgery were not included in analyses. Recordings were continuous and stored in 1-min bins.

### Ovariectomy and Silastic Capsule Replacement

Ovariectomies were performed at weaning (p21) at the same time as the implantation of the E-Mitter, as previously described(10, 57). The E-Mitter surgery served as a control operation in non-OVX animals. Incisions were closed using dissolvable sutures and wound clips. At p29, OVX animals were anesthetized and implanted with silastic capsules (0.78 mm I.D., 1.25 O.D.; Dow Corning, Midland, M). Capsules were implanted subcutaneously and intrascapular. Capsules were 20mm in length with with 5mm silicone sealant at each end (Sigma Aldrich, St. Louis, MO) and contained either 112µg (180µg/mL) 17β estradiol in sesame oil, or sesame oil alone. E2 treatment results in plasma E2 concentrations averaging ∼ 5µg/day, beginning in animals of 80-100g(110, 111). Capsules were primed for 24 h prior to implantation via submersion in 0.9% saline in order to avoid delivering a large initial bolus of E2. Although these doses have been tested previously, large variability in serum E2 levels following silastic implant is typically reported(110–113). Incisions were closed using dissolvable suture and a wound clip, and buprenorphine was delivered as above for post-operative analgesia.

### Contraceptive Administration

Ethinyl Estradiol (EE2, 30µg/day) and Levonorgestrel (30 µg/day), a progestin, were dissolved in in 0.01 mL of sesame oil and delivered subcutaneously at the nape of the neck daily for the duration of two estrous cycles during mid to late adolescence (p50 to p58). Although a wide range of rodent doses of EE2 and Levonorgestrel have been reported(114–116), the doses chosen here aimed to match those used consistently for suppressing ovulation and mimicking effects observed in humans, such as increased blood pressure(117), and for comparability to existing rodent literature on subcutaneous delivery of Levonorgestrel and EE2(118–120). Many doses for orally delivered EE2 and Levonorgestrel in rats also fall in this range(82,115,121,122). These drugs have been available as human contraceptives for decades under several brand names in widely varying doses(100, 123).

### Fecal Sample Collection

Fecal E2 (fE2) concentrations were assessed across puberty from feces generated over 24 h periods. Fecal samples provide hormone concentrations more representative of average daily hormone concentrations than single timepoint blood samples(124–128) and eliminates associated stress and infeasibility of high-frequency, longitudinal blood collection. Samples were collected in small, airtight bags at the end of dark phase under dim red light (<5 lux) from p25 to p37 (pre puberty and first cycle), p45 to p51 (mid-puberty), and p55 to p65 (late puberty to early adulthood) in all groups, and additionally to p75 in Intact+C and Intact groups (adulthood). Samples soiled with urine were discarded, and all other droppings generated over each 24-h segment were combined. Within 1 h of collection, samples were stored at -20°C until processed. Sample collection was rapid (∼ 1 min per animal).

Before assessment of hormone concentration, samples were processed according to manufacturers’ instructions(129). Briefly, samples were placed in a tin weigh boat and heated at 65°C for 90 minutes, until completely dry. Dry samples were ground to a fine powder in a coffee grinder, which was wiped down with ethanol and dried between samples to avoid cross contamination. Powder was weighed into 0.2 mg aliquots. For hormone extraction, 1.8mL of 100% ethanol was added to each test tube, and tubes were shaken vigorously for 30 minutes. Tubes were then centrifuged at 5,000 RPM for 15 minutes at 4°C. Supernatant was moved to a new tube and evaporated under 65°C until dry (∼ 90 minutes). Sample residue was reconstituted in 100uL of 100% ethanol. 25µL of this solution was diluted for use in the assay and remaining sample was diluted and stored.

### ELISA Assays

A commercially available fE2 enzyme-linked immunosorbent assay (ELISA) kit was used to quantify E2 in fecal samples (Arbor Assays, Ann Arbor, MI). These assays have been previously published in species ranging from rats and mice(130–132), to wolves(133), to humans(134). ELISAs were conducted according to manufacturer’s instructions. To ensure each sample contained ≤ 5% alcohol, 25µL of concentrate were vortexed in 475µL assay buffer. All samples were run in duplicate, and an inter-assay control was run with each plate. Sensitivity for the assay was 39.6 pg/mL and the limit of detection was 26.5 pg/mL. fE2 intra-assay coefficient of variation (COV) was 5.94% and inter-assay COV was 5.71%.

### Data Availability and analysis

All code and data used in this paper are available at A.G.’s and L.J.K.’s Github Respository (https://github.com/azuredominique; https://github.com/Kriegsfeld-Lab)(135). Code was written in MATLAB 2020b with Wavelet Transform (WT) code modified from the Jlab toolbox and from Dr. Tanya Leise(136, 137). Briefly, data were imported to MATLAB at 1-min resolution. Any data points outside ± 4 standard deviations were set to the median value of the prior hour, and any points showing near instantaneous change, as defined by local abs(derivative) > 10^5^ as an arbitrary cutoff, were also set to the median value of the previous hour. Small data gaps resulting from intermittent data collection (<10 minutes) were linearly interpolated. Continuous data from p26 to p74 were divided into three equal-length phases: early to mid puberty (p26 to p41), mid to late puberty (p42 to p58), and late puberty to early adulthood (p59 to p74).

### Wavelet Analyses and Statistics of CBT Data

Wavelet Transformation (WT) was used to generate a power estimate, representing amplitude and stability of oscillation at a given periodicity, within a signal at each moment in time. Whereas Fourier transforms allow transformation of a signal into frequency space without temporal position (i.e., using sine wave components with infinite length), wavelets are constructed with amplitude diminishing to 0 in both directions from center. This property permits frequency strength calculation at a given position. In the present analyses we use a Morse wavelet with a low number of oscillations (defined by β=5 and γ=3, the frequencies of the two waves superimposed to create the wavelet(138)), similar to wavelets used in many circadian and ultradian applications(9,10,136–139). Additional values of β (3–8) and γ (2–5) did not alter the findings (data not shown). As WTs exhibit artifacts at the edges of the data being transformed, only the WT from p26 to p74 were analyzed further. Periods of 1 to 39 h were assessed. For quantification of spectral differences, WT spectra were isolated in bands; circadian periodicity power was defined as the max power per minute within the 23 to 25 h band; ultradian periodicity power was defined as the max power per minute in the 1 to 3 h band. The latter band was chosen because this band corresponded with the daily ultradian peak power observed in ultradian rhythms (URs) across physiological systems in rats(6,140–142).

For statistical comparisons of any two groups, Mann Whitney U (MW) rank sum tests were used to avoid assumptions of normality for any distribution. Non-parametric Kruskal-Wallis tests were used instead of ANOVAs for the same reason; for all Kruskal-Wallis (KW) tests, χ^2^ and *p* values are listed. In cases of repeated measures within an individual over adolescence, Friedman’s tests were used. Data from each estrous cycle were treated as independent. We chose to treat estrous cycles as independent for 3 reasons: 1) the estrous cycle is the longest periodicity rhythm within the study, 2) the adolescent estrous cycle develops rapidly from one iteration to the next in rats, and 3) because the interventions of ovariectomy and birth control administration exert their effects by removal or modification of estrous cycles. Dunn’s test was used for multiple comparisons. Mann Kendall (MK) tests were used to assess trends over time in wavelet power and linear CBT over three equally sized temporal windows described above. For short term (<3 days of data) statistical comparisons, 1 data point per hour was used; for longer term (>3 days of data) statistical comparisons, 1 data point per day was used. Continuous wavelet power data were smoothed with a 24 h window using the MATLAB function “movmean”. Violin plots, which are similar to box plots with probability density of finding different values represented by width(143), were calculated using the MATLAB function “violin”. Median daily circadian power was regressed against each day’s fE2 for each group using a mixed effects linear regression (MATLAB function “fitlme”, formula: circadian daily medians∼1+E2 values + ( 1+ E2 |Individual ID).

### Data alignment and Analysis of E2 Concentrations and Estrous Cyclicity

The day of fE2 rise before cycling began was defined as the first day fE2 level rose > 2 standard deviations above its starting value at p25. The initial rise in fE2 was used as an alignment point for CBT, ultradian power, and Z-Score(CBT) - Z-Score(UR power) group averages. Group differences in fE2 area under the curve by cycle were assessed using the MATLAB function “trapz” and KW tests with Dunn’s post hoc correction. As estrous cycles are not all aligned in time or by age, samples were aligned with the highest value in a collection period (e.g., mid puberty) where a ‘fall’ was observed three days later. Fast Fourier Transforms (FFT) were used to assess the presence or absence of 4-5 day power in CBT in each individual from the period of fE2 rise until p50 (when Intact+C animals started receiving daily contraceptive injections), and from p50 to p74. In order to further assess commencement and stability of estrous cycling after first rise in fE2, as well as any potential perturbation during and after contraceptive administration, metrics were divided into 4 day blocks, with each day labelled 1,2,3, and 4: repeating for subsequent cycle lengths. Groups for statistical comparison were constructed from all data corresponding to 1’s, 2’s, 3’s and 4’s. Friedman’s tests with Dunn’s correction for multiple comparisons were used to determine if values associated with each day of cycle (e.g., all day 1’s) varied significantly from other days of the cycle by group.

## Results

### Impact of Hormonal Status on Estradiol Concentrations and Weight Gain Across Adolescence

Frequent fecal estradiol (fE2) measurements were collected to assess if hormonal status affected the level or temporal patterning of fE2 across puberty. FE2 concentrations did not differ between groups from p25-p31, a baseline period prior to puberty onset (χ^2^=4.48, p=0.214; **Figure 1A**). Vaginal opening occurred between p31 and p33 in Intact rats, and fE2 rose 2 standard deviations above its p25 starting value between p31 and p36.

**Figure 1.**
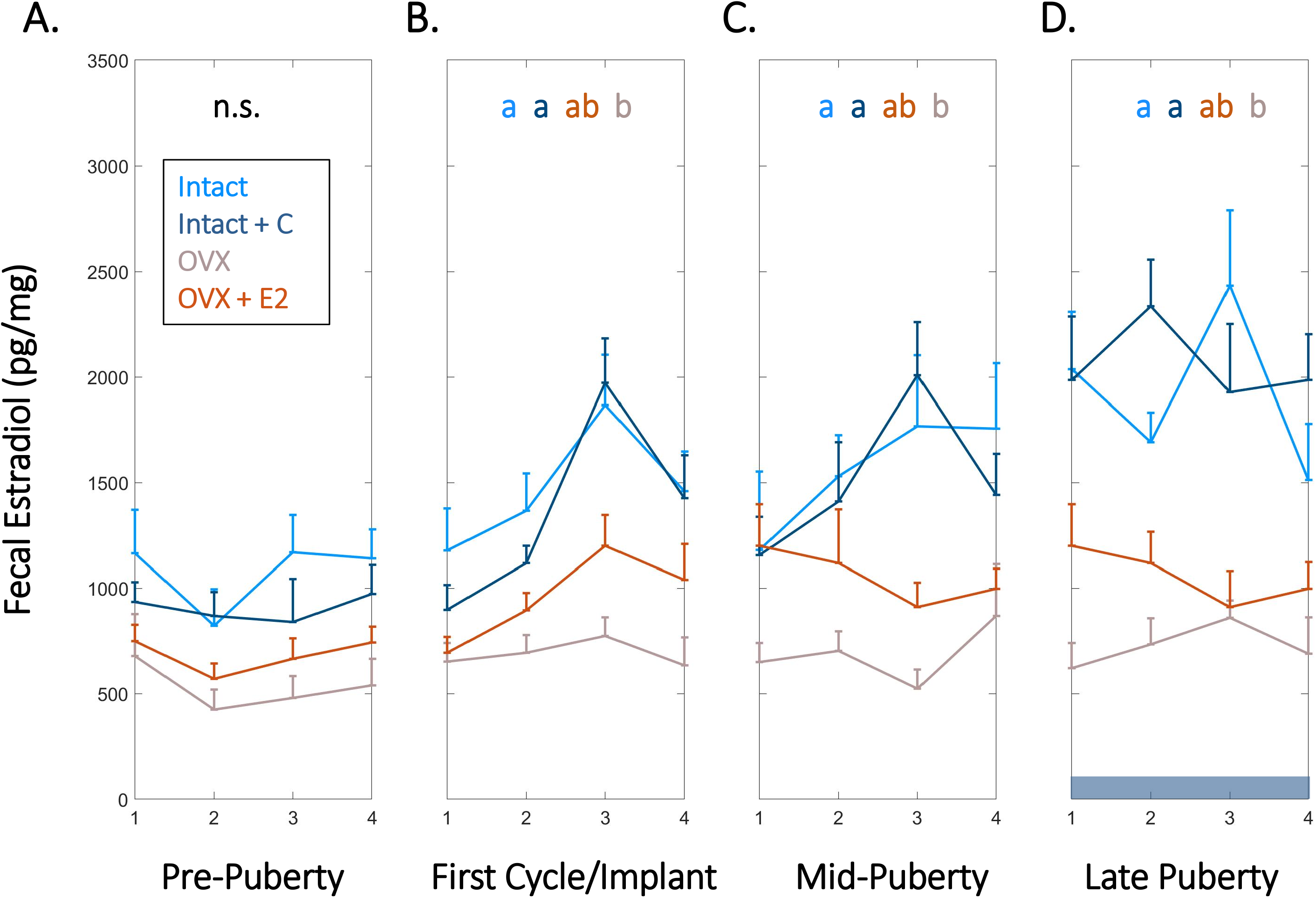
High Frequency Fecal Estradiol Enables Monitoring of Estrous Cycle Emergence, Sex Steroid Manipulation, and Ovarian Status During Adolescence. Group mean (+S.E.M.) fecal estradiol concentrations of Intact (light blue), intact + short-term pubertal contraceptives (Intact+C; dark blue), OVX (gray), and OVX+E2 (orange) groups did not significantly differ prior to puberty (p24-30) (A). Fecal estradiol in Intact and Intact+C groups increased over that of OVX animals beginning at the first cycle following vaginal opening or relative to the time of silastic implant in OVX+E2 animals (p30-37) (B) and remained significantly elevated thereafter at mid puberty (p43-p49; C) and during late puberty (p55-61; D). Dark horizontal bar in D indicates Intact+C data were gathered during contraceptive administration. Color of letters at the top of B, C, and D indicate experimental group; letters indicate statistical differences, with groups not sharing the same letter being significantly different (p<0.03).

During this window, Intact and Intact+C animals’ fE2 concentrations exceeded that of OVX animals (χ^2^=15.9, p=0.001; p=0.0134 and p=0.003, respectively; **Figure 1B**). This difference was maintained at mid puberty (cycles aligned from p40 to p47) (χ^2^=13.7, p=0.003; p=0.003 and 0.032, respectively) and early adulthood (cycles aligned from p55 to p61) (χ^2^=17.1, p=0.001; p=0.001 and 0.009, respectively; **Figure 1C-D**). OVX+E2 animals were not different from other groups at any timepoint, with intermediate values between Intact and OVX groups (p>0.05 in all cases). Unlike Intact animals, Intact+C animals did not exhibit days of elevated fE2 every 4^th^ day (See **Supplemental Figure 1**). However, fE2 concentrations did not differ between Intact and Intact+C groups approximately 4 to 5 cycles after contraceptive administration ceased, between p69 to p75 (χ^2^=1.62 p=0.203). See **Methods** for details of within-cycle alignment.

Additionally, daily weights were measured to recapitulate known effects of E2 on pubertal growth trajectory, and to assess if contraceptive administration modulated weight gain. Intact animals gained weight consistently across puberty. Pre-pubertal OVX was associated with increased body weight at mid puberty, with OVX animals weighing more than animals from all other groups, and OVX+E2 animals weighing more than Intact or Intact+C groups (χ^2^=57.9, p=1.65*10^-12^; p<0.05 for all individual comparisons). By early adulthood, both OVX and OVX+E2 groups weighed significantly more than Intact or Intact+C groups and did not differ from one another (χ^2^=76.1, p=2.08*10^-16^; p=0.99 for OVX vs. OVX+E2; p<0.05 for all other comparisons). Eight days of contraceptive administration did not significantly impact weight relative to Intact animals (χ^2^=0.28, p=0.594; See **Supplemental Figure 2**).

### Circadian, but not Ultradian, Power of Body Temperature Increased Across Pre-to-Mid Adolescence

As reproductive circadian and ultradian rhythms change markedly across adolescence and may be coupled to CBT(32,49,144), we investigated the impact of estradiol status on the timing and tempo of CBT rhythmicity. All animals exhibited a significant positive trend in CR power from pre-to-mid adolescence (p26 to p41) (p=0.002, 0.007, 0.029, 0.04 for Intact, Intact+C, OVX, and OVX+E2 animals, respectively). CR power stabilized thereafter (p>0.05 in all cases; **Figure 2A-B**). To examine the relative rate of this increase across groups, we set a criterion of 2 standard deviations above the mean. CR power rose 2 standard deviations above the mean significantly faster in OVX+E2 animals compared to Intact or OVX animals (χ^2^ =19.0, p=3*10^-4^; p=0.025 and 0.001, respectively; **Figure 2L**).

**Figure 2.**
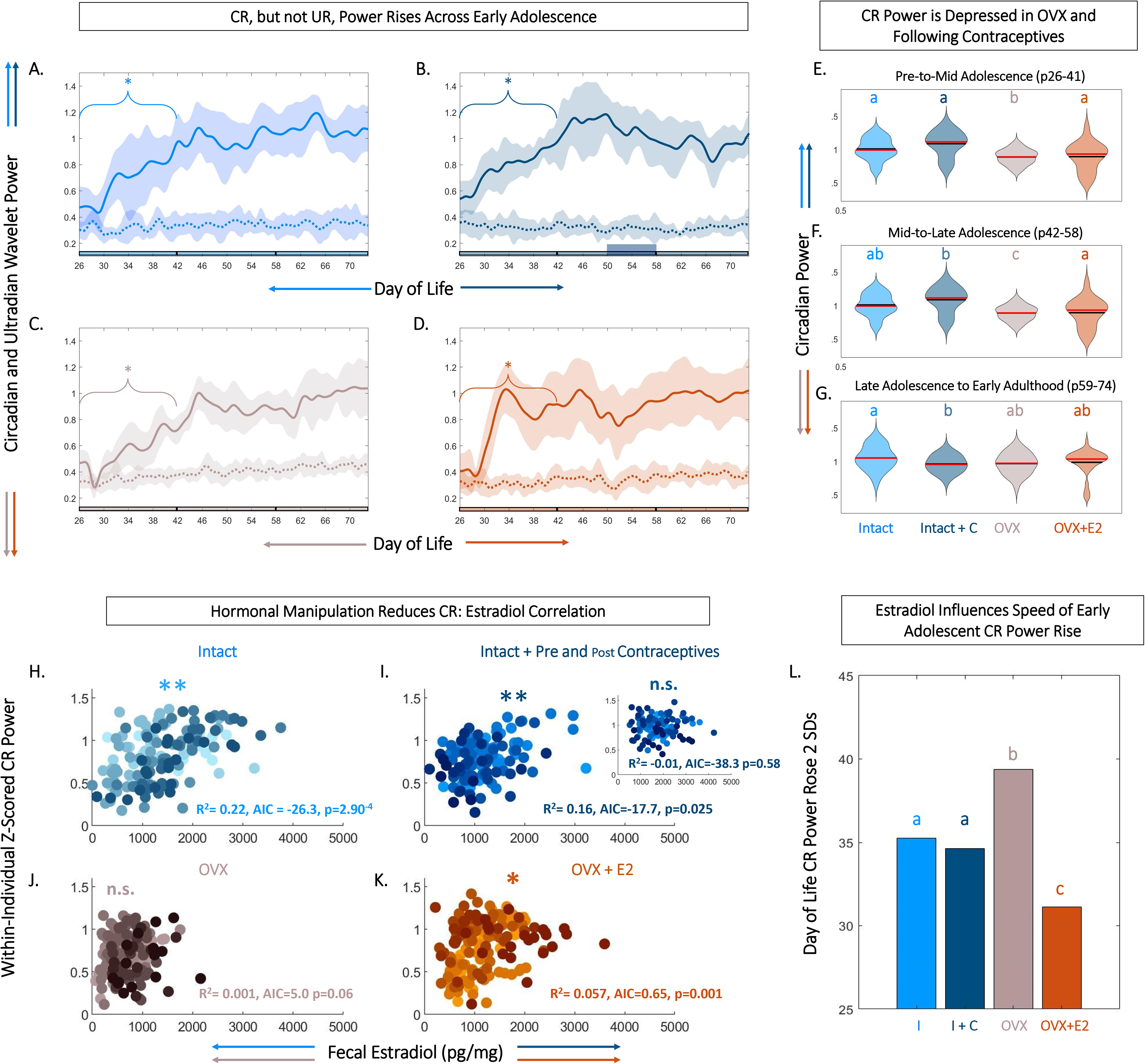
Level and Rate of Early Adolescent Rise in CBT Circadian Power are Tied to Estradiol. Circadian, but not ultradian power rises across early adolescence (A-D). Linear plots of group mean (± S.E.M.) of CBT circadian (solid) and ultradian (dashed) power in Intact (light blue, A), Intact+C (dark blue, B), OVX (gray, C) and OVX+E2 (orange, D) animals. * Indicates significant trend over time for the bracketed time region (p<0.05). Phase of adolescent time periods (pre to mid, mid to late, and late to adult) are indicated by black dividers in the colored x-axis at p42 and p58. Although CBT circadian power rises over early adolescence in all groups, estradiol increases the rate of this rise (L). Violin plots (E-G) of circadian power in each group analyzed by segment of life: pre to mid adolescence from p26-41 (E), mid to late adolescence from p42-57 (F), and late adolescence through early adulthood from p58-73 (G) illustrate that CR power is highest in Intact animals, with reduction following hormonal contraceptive administration and OVX, with a partial rescue in OVX+E2 animals. Black lines indicate mean and red lines indicate median of each plot. Scatters of fE2 level by CR power indicate that hormonal manipulation reduces or eliminates the correlation between CR power and fE2 concentrations (H-K). Note that each individual within a group is plotted in a unique color. CR power and fE2 are significantly correlated in Intact (light blue, H), Intact+C animals prior to contraceptive administration (dark blue, I), with inset depicting abolished correlation during and after contraceptive administration. CR power and fE2 are weakly correlated in OVX+E2 (orange, K) but not OVX animals (J). * Indicates significant positive correlation between fE2 and circadian power. AIC indicates relative performance of the mixed effects model. Color of letters at the top in E-G and L indicate experimental group; letters indicate statistical differences, with groups not sharing the same letter being significantly different (p<0.03).

Ultradian power did not show a significant upward or downward trend across the study period in any group (p>0.05 for all groups at all time windows) (**Figure 2A-D**). Although directionality of CR power change across adolescence was similar (i.e., an early increase followed by a plateau) in all individuals, magnitude of circadian power differed among groups. Specifically, circadian power for OVX animals was depressed compared to all other groups from pre to mid adolescence (χ^2^ =125, p=4.02*10^-27^, p<0.02 for OVX vs. all other groups; **Figure 2E**). From mid to late adolescence (p42 to p58), CR OVX power remained depressed and OVX+E2 trended toward lower power (χ^2^ =112, p=2.82*10^-24^; p<0.01 Intact and Intact+C vs OVX; **Figure 2F**). In early adulthood (p59 to p74), following contraceptive administration, the CR power for Intact+C was depressed compared to Intact rats (χ^2^=37.9, p=2.93*10^-0.8^, p=0.04; **Figure 2G**).

### Body Temperature Circadian Power was Most Correlated to Fecal Estradiol Level in Unmanipulated Animals

Temperature level, UR power, and E2 exhibited coupled patterning, and appeared to change markedly during adolescence. However, it was unclear if CBT circadian rhythmicity was coupled to estradiol, if such a relationship existed during adolescence, and if the relationship could be modified by hormonal state. FE2 and normalized circadian power were strongly correlated in Intact (R^2^=0.226, p=2.90*10^-4^) and Intact+C rats prior to p50 when contraceptive administration began (R^2^=0.166, p=0.025), and weakly correlated in OVX+E2 animals (R^2^=0.062, p=0.001; Figure 2 H-K). This positive correlation was abolished during and after hormonal contraceptive administration in the Intact+C group (R^2^=1.06*10^-5^, p=0.58; **Figure 2I, inset**). Circadian power and fE2 was not significantly correlated in OVX animals (R^2^=0.013, p=0.06; **Figure 2J**).

### Ovulatory Rhythms in CBT and Perturbations During and After Contraceptive Administration

We next investigated the relationship between ORs and cycles in body temperature. Our goal was to determine if interactions at this timescale a) could be observed in continuous body temperature in adolescents, b) if previously described hormonal UR modulations by phase of cycle(145, 146) translates to ovulatory cycles in CBT URs, and c) if exogenous hormone administration disrupts these continuous dynamics. In intact rats, we observed a significant 4-day modulation of combined CBT and UR power corresponding to the estrous cycle (Intact χ^2^ =59.1, p=9.26*10^-13^, Intact+C group prior to BC administration χ^2^ =13.7 p=0.003; **Figure 3A-B, 4A-B**). This 4-day pattern commenced in Intact and Intact+C rats (prior to treatment) with a significant increase in mean daily CBT following the first rise in fE2 above 2 standard deviations (p=0.03 in each case). Intact and Intact+C (prior to treatment) animals also exhibited a 4-day pattern of UR power modulation. This modulation manifested as as a significant trough of UR power within 4 days of the first rise of fE2, as previously reported in adult rodents(9,10,147,148) (p=0.04, p=0.03, respectively). The combination of UR power and linear temperature yielded a more easily separable metric which rose significantly the day after first rise of fE2 in Intact and Intact+C (p=0.01, p=0.02, respectively; **Figure 3A-B**). (For individual metric comparisons see Supplemental Figure 3).

**Figure 3.**
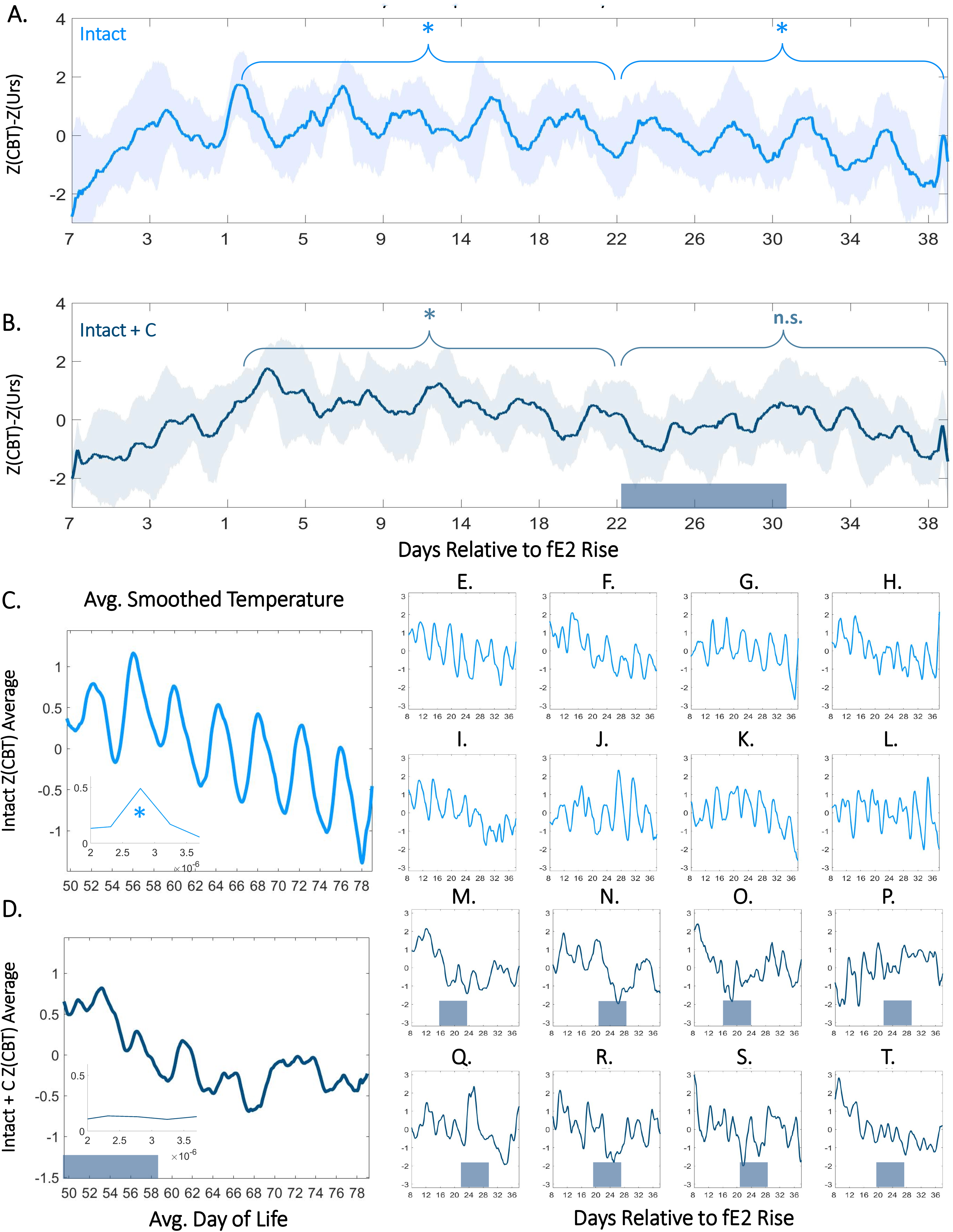
Contraceptive Administration in Adolescence Persistently Perturbs 4-Day Temperature Rhythms. Normalized CBT mean (± S.E.M.) minus UR power relative to within-individual first day of fE2 rise in Intact (A) and Intact+C (B) rats (see: Methods and Supplemental Figure 3). Dark bars along the x-axis for Intact+C animals indicate average time of contraceptive administration relative to fE2 rise. * Indicates regions of time over which every 4^th^ day’s CBT values are significantly elevated compared to other days of cycle (p<0.003). Twenty-four hour smoothed average plots of normalized linear CBT in Intact (C) and Intact+C (D) individuals from the time of contraceptive administration illustrate a reduction in regularity of oscillations. Insets show FFT centered at 4-5 days. * Indicates significantly higher AUCs in the 4-5 day range for Intact (panel C) compared to Intact+C rats (panel D). Individual animals (E-T) comprising Intact (light blue) and Intact+C (dark blue) groups prior to and following hormonal contraceptive administration. Dark bars along horizontal axes indicate time of contraceptive administration; administration days differ based on an individual’s day of fE2 rise.

We noted an absence of significant 4-day differences in combined CBT and UR power in the Intact+C group during hormonal contraceptive administration, even following 4 cycle-lengths of recovery (Intact+C group χ^2^ =7.2, p=0.07; Intact group over same time period χ^2^ =58.9, p=1.00*10^-12^). A FFT of data in Intact and Intact+C animals prior to contraceptive administration revealed comparable AUCs for 4 to 5 day oscillations (no group difference; χ^2^ =0.54, p=0.46; **Figure 3C**, **Inset**). However, after hormonal contraceptive administration, AUC for Intact animals was significantly greater for 4 to 5 day oscillations than in Intact+C animals (χ^2^ =3.98, p=0.046; **Supplemental Figure 4**). As expected, a 4-day pattern was also absent in OVX and OVX+E2 animals (p>0.05 in both cases). Note that 5-day cycles occurred rarely in Intact rats and using 5-day bins rather than 4-day bins abolished significant differences by day of cycle for all groups (*data not shown*).

### CBT Increased in Early to Mid Adolescence

As growth and metabolism speed in adolescence, we hypothesized that CBT would also increase during the period of most rapid growth, pre to mid puberty. Pre to mid puberty (p26-41) was associated with a significant positive trend in CBT in Intact (p=5.75*10^-5^) and Intact+C animals prior to contraceptive administration (p=1.20*10^-4^; **Figure 4A-B**). Notably, implantation of the silastic capsule in OVX and OVX+E2 animals resulted in a transient (one day) increase in CBT (OVX p=0.01, OVX+E2 p=0.03; **Figure 4C-D, Supplemental Figure 5**). This surgical-recovery-associated rise was highly variable and did not differ between OVX and OVX+E2 animals (p=0.65; **Supplemental Figure 5**). Interestingly, the early pubertal CBT increase did not require E2, as OVX animals also exhibited a significant positive trend (p=0.034; **Figure 4C,E**). For a summary guide to CBT features that may be useful for Intact pubertal staging, see **Supplemental Figure 6**).

**Figure 4.**
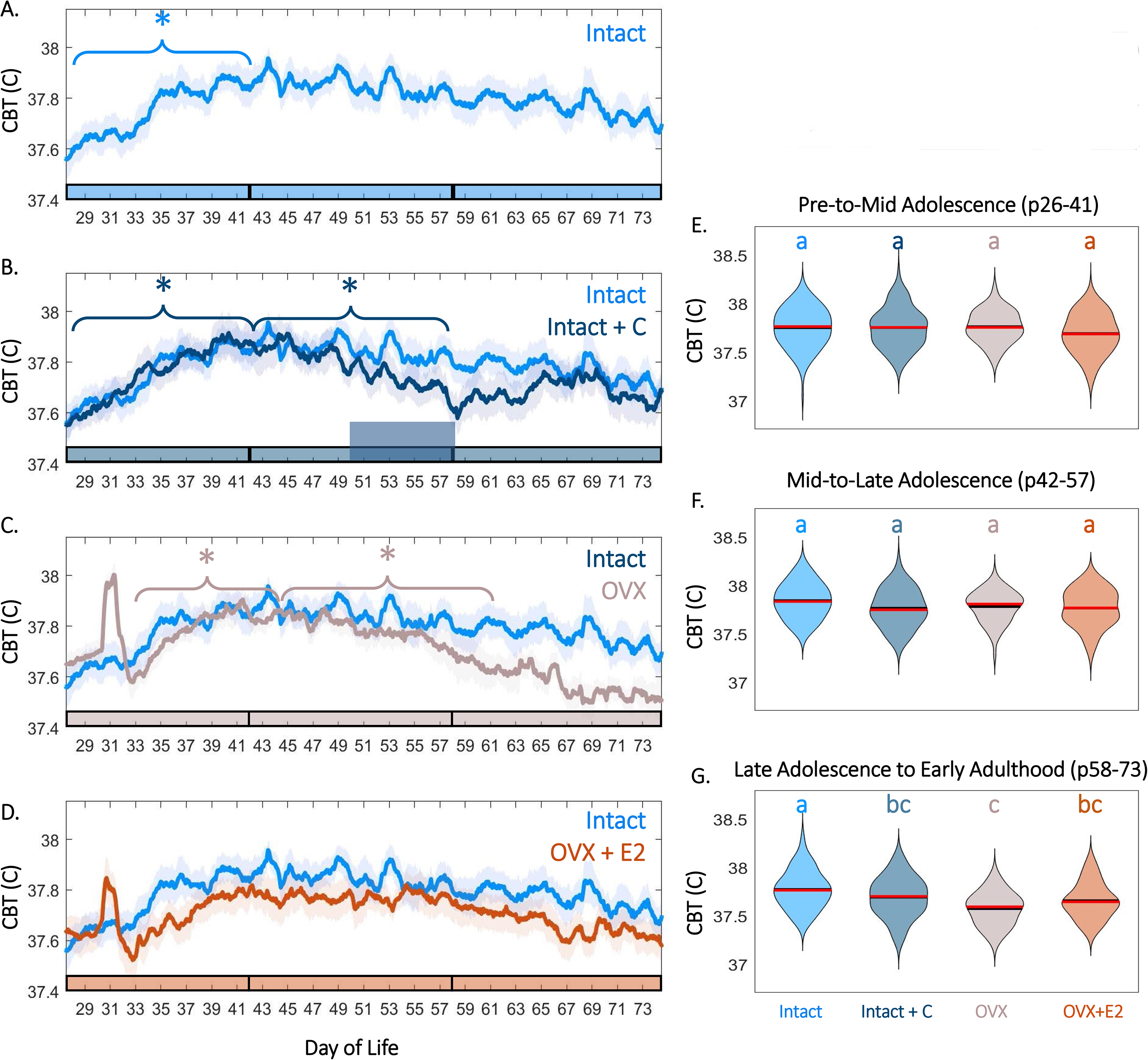
Female Adolescence is Associated with Sex Steroid-Dependent CBT Levels and Trends. CBT linear group means (± S.E.M.) in Intact (A, light blue) compared to Intact+C (B, dark blue), OVX + sham (C, gray), and OVX+E2 (D, orange) animals. * Indicates significant trend during the bracketed time period for the group matching the color of the bracket (p<0.03). Phase of adolescence time periods (pre to mid, mid to late, and late to adult) are indicated by breaks in the colored x-axis at p42 and p58. Time of hormonal contraceptive administration in panel B is indicated by the tall horizontal bar. Violin plots of temperature for all groups at early to mid adolescence (E), mid to late adolescence (F), and early adulthood (G) indicate that hormonal contraceptive administration leads to reductions in CBT relative to controls after administration. Ovariectomy, even with E2 replacement, is also associated with significantly reduced temperatures by early adulthood (G). Color of letters at the top in E-G indicate experimental group. Letters indicate statistical differences, with groups of different letters being significantly different (p<0.001).

### CBT Maintenance in Late Adolescence to Adulthood Required Estradiol

Complex interactions exist between metabolism, growth, and E2 level during adolescence. As estrogen deficiency in puberty is associated with weight gain and reduced metabolism, we investigated if maintenance of elevated temperature would be impacted by hormonal status. The maintenance of increased temperature in late puberty and adulthood was E2-dependent, with OVX animals exhibiting a significant downward trend in CBT from mid to late puberty (p58-p74; p=0.01) relative to Intact and OVX+E2 animals (p>0.05 in each case; **Figure 4C-D**). E2 treatment in the OVX+E2 group prevented intra-individual CBT decline in late puberty; correspondingly, temperatures in the OVX but not OVX + E2 groups were lower than that of Intact animals (late puberty to early adulthood χ^2^ =62.8 p=1.46*10-^13^, p=4*10^-4^ for Intact vs. OVX, p=0.16 for Intact vs. OVX+ E2; **Figure 4F-G, Supplemental Figure 5**).

### Contraceptive Administration Longitudinally Depressed CBT

CBT power did not exhibit a positive or negative trend from mid puberty through early adulthood (p42 to p58) in Intact animals (p=0.12), but exhibited a significant downward trend in Intact+C animals during the period of contraceptive administration (p=0.028; **Figure 4A-B**), resulting in a trend toward depressed temperatures following administration in mid to late adolescence (χ^2^=21.84 p=7.04*10^-05^, p=0.1 for Intact vs. Intact+C) that persisted into early adulthood (χ^2^=62.83 p=1.46*10^-13^, p=0.1 for Intact vs. Intact+C; **Figure 4B, 4F-G**).

## Discussion

### Adolescent Development and Estradiol Dependence of CBT Rhythmicity

Female adolescence is characterized by stereotyped development of CBT rhythms at the ultradian, circadian, and ovulatory timescales. The present findings reveal that early adolescence in the female rat is marked by rising CBT and CBT circadian power, and commencement of 4-day cycling in CBT and CBT URs. These early circadian and ultradian changes likely reflect maturation of the reproductive axis, as 1) the rate of CR power rise was hastened by estradiol, 2) the commencement of 4-day temperature cycling was preceded by a rise in fE2, and 3) CBT CR power was correlated with fE2 concentration(65, 149). These observations confirm and extend reports of early pubertal development of ultradian-circadian-ovulatory interactions(46) (47), and suggest that reproductive-thermoregulatory coupling may develop prior to, or in tandem with, pubertal onset.

Adolescent increases in UR amplitude for many endocrine outputs are well-documented, but UR structure across all of puberty is not well mapped(28, 65). However, from the time of rise in fE2 (a defining marker of pubertal onset), CBT UR power retained the same mean value but commenced a 4-day, ovulatory cycle-associated pattern. Four day cycles in URs are consistent with data collected on URs in adult rodents(10,147,148) and humans (at a longer time scale)(139). Four-day patterning was not present in ovariectomized or E2-replaced animals, consistent with dependence on the ovarian cycle. These results support that URs in CBT achieve adult amplitude and stability early in life, prior to maturity of CRs(150–152), and that interactions among thermoregulatory and reproductive circuits permitting rapid *ultradian* coupling are established well before pubertal onset.

Additionally, the present findings suggest that some dynamics of CBT development require specific patterns of E2 rather than simply concentrations above a particular threshold. Ovariectomy eliminated ORs, increased weight, reduced CR power, rate of CR power rise, and correlation between CR power and fE2, and overall temperature(153, 154). E2 replacement partially rescued circadian metrics and reduced weight but did not recapitulate ORs. Short-term exposure to contraceptives in late adolescence longitudinally altered CBT metrics by reducing temperature and CR power, and abolishing ORs in CBT and UR power. Despite the apparently early maturation of substrate for thermoregulatory and reproductive coupling, the impact of E2 replacement and the enduring effects of short-term contraceptives suggest that these systems are sensitive to both level and patterning of reproductive hormones across the adolescent period.

Together, temperature amplitude and oscillation stability at the UR, CR, and OR timescales are modulated across the adolescent transition and are influenced by endogenous and exogenous E2. CBT URs mature to their adult amplitude and stability prior to CRs, and ORs and are rapidly modulated by ovulatory phase in intact animals (**See Supplemental Figure 6**). Conversely, CRs increase in magnitude and stability in early adolescence, are not significantly impacted by the phase of OR, and are impacted by ovariectomy. These results support the notion that ultradian and ovulatory systems are tightly coupled and reflected in CBT, and that circadian effects on the ovulatory cycle may be unidirectional(22, 139). Ovariectomy, E2 replacement, and pubertal contraceptive administration have a number of effects on rhythmic dynamics at each timescale, overall indicating that ‘intact’ dynamics are not easily recapitulated and that exogenous sex steroids can have enduring impact. *Limitations*. The ovulatory cycle of the female rat differs from that of humans, in that rats do not exhibit prolonged elevation of progesterone in the absence of pregnancy or pseudopregnancy(155). This lack of a true luteal phase means that post-ovulatory temperature elevation in the rat follows a more compressed trajectory than in humans(156). Therefore, exposure to estrogen and progesterone analogs in rats for prolonged periods may be associated with a different phenotype in rats than in humans(157). Finally, the laboratory environment imposes artificially stable environmental conditions on animals; recapitulation of these patterns under naturalistic conditions in future experiments will strengthen the translational potential of this work.

### Considerations of Rhythmicity Perturbation Through Adolescent Contraceptive Use

Although rats cannot fully model human biology, it was notable that contraceptive administration imposed lasting structural changes on CBT rhythmicity and its relationship with E2. Levonorgestrel and EE2 administration abolished the 4-day modulation of fE2, UR power, and temperature level, eliminated the correlation between CR power and fE2, and significantly depressed CR power. As these changes required continuous monitoring to detect and occurred in the absence of significant group changes to fE2 level, it is not unreasonable to speculate that studies of less frequently timed samples, or samples averaged across individuals, could make similar disruptions in humans difficult to detect.

Perturbation of body temperature rhythms is associated with diverse health insults across species, both reflecting perturbation in underlying systems and potentially acting as a causative agent. Circadian disruption to CBT rhythms occurs in, and is proportional to, severity of jetlag(158), depression(159), sepsis severity(160, 161), post-traumatic injury(162), cognitive decline(163), and has even been proposed as a root cause of disease dubbed “Circadian Syndrome”(164) . Disruption to ovulatory temperature rhythms occurs in anovulatory and atypical luteal phase cycles(165–167), including those arising from polycystic ovarian syndrome (PCOS)(168). The impact of ultradian rhythmic disruption of the reproductive axis requires additional study(169–171), but existing work in other hormonal systems suggests that preservation of pulsatility in drug delivery (e.g., of cortisol in Addison’s disease or insulin in diabetes) can lead to better patient outcomes when compared to conventional non-rhythmic treatment(66,67,104,172,173). It is likely that disruption of URs may result in negative impact analogous to disruption at longer timescales. It may appear counter-intuitive that thermoregulation could be both a reporter for such diverse maladies and a potential mediator for disease progression. However, temperature rhythm disruption is associated with a wide range of temporal, inflammatory, and endocrine insults, in part, because thermoregulatory circuits are directly impacted by the master clock, and modulated by inflammatory factors, autonomic status, and a variety of endocrine factors including estradiol(8, 174) and progesterone(7). Lastly, CBT itself acts as a synchronizing cue for peripheral circadian oscilllators(175). Further research is needed to disentangle if disruption of CBT rhythmicity itself causes harm, or if CBT rhythms are merely reporting perturbations in underlying systems (e.g., sex hormones or metabolic factors).

Our observations suggest that a non-physiological pattern of contraceptive administration may act as ‘hormonal jetlag’. The degree of lasting impact to other systems that rely on temperature as an entraining stimulus, or disrupted systems reported indirectly by temperature (e.g., SCN, endocrine, autonomic) remain to be assessed. Furthermore, the extent to which such disruptive effects differ among species(157), contraceptive agents, administration methods, or between adolescent populations and adults requires further investigation. Encouragingly, the observation that rodents and humans exhibit similar CBT patterning during the peri-ovulatory period(10,139,140) points to potential translational relevance

Together, despite established societal benefits of widely available hormonal contraception(176), especially in individuals experiencing hormonal irregularities(177), the present findings suggest that administration of exogenous estrogens and progestins during adolescence leads to persistent rhythmic disruption across timescales. Future research is needed to determine if rhythmic patterns of sex steroid administration more closely mimicking endogenous release, analogous to those implemented in cortisol(67) and closed loop insulin therapy(178), can minimize rhythmic disruption. Conversely, future studies that validate an “updated” symptom-thermal method using signal processing of continuous CBT data may provide feasible, non-disruptive alternatives for contraception(106,139,179–181). Contraceptive administration to adolescent girls is on the rise(74), and there are a paucity of data on the impact of chronic hormonal perturbation on endogenous rhythmicity, or if such disruptions during the sensitive window of adolescence have lasting effects(93,94,101,182,183). Further research is needed to characterize the impact of adolescent hormonal contraceptive use so that further improvements can be made, individuals at-risk for side effects can be identified, and informed decisions about family planning can be made.

### Utility of CBT for Monitoring Pubertal Development in Rodents and Potential Translational Relevance

Continuous monitoring of CBT may have great utility for passive detection of pubertal milestones in rodents in preclinical research. Existing methods for pubertal staging in rodents carry considerable downsides: frequent blood sampling and vaginal lavage(184) are repeatedly invasive, and fecal hormone analysis is time consuming and costly(124, 185). Moreover, our results indicate that individual rats do not traverse identical pubertal trajectories by day of life, and that this assumption could lead to considerable errors in staging. Conversely, signal processing of passively collected CBT can add temporal resolution, greater quantitative power, and limit repeated invasive procedures for staging puberty(186).

In human subjects, continuous temperature monitoring via wearables may similarly facilitate the study of pubertal development. The signal characteristics of rat CBT exhibit remarkable similarities to human peripheral temperature (See:(61,148,187)). Monitoring human peripheral temperature during adolescence could serve broad purposes – from personalizing health education based on self-collected data(188, 189), to adapting teaching style to an individual’s developmental phase(190–193), to enabling research into the impact of teen contraceptive use(194, 195) or the process of gender transition(196, 197). Together, pubertal monitoring via continuous body temperature is worthy of further investigation in both animal models and human subject populations for its potential utility to individuals, researchers, families, and clinicians.

## Conclusion

In conclusion, body temperature monitoring provides a window into the development of biological rhythms in puberty over multiple timescales. These rhythms may serve as convenient and high temporal resolution indicators of developmental stage and trajectory for application in research and clinical studies. Our study of body temperature also reveals unintended side effects of tonic hormonal manipulations. We anticipate that these findings will inform creative improvements to female reproductive research and healthcare.

## Acknowledgements

The authors would like to thank Andrew Ahn, Ronald Dahl, Frederic Théunissén, and Albert Qü for their helpful feedback on study design and methods and invaluable comments and suggestions on an earlier version of this manuscript.

## Funding

This work was funded by SL-CN grant #1640885, NIH grant HD-050470

---

**Supplemental Figure 1.**
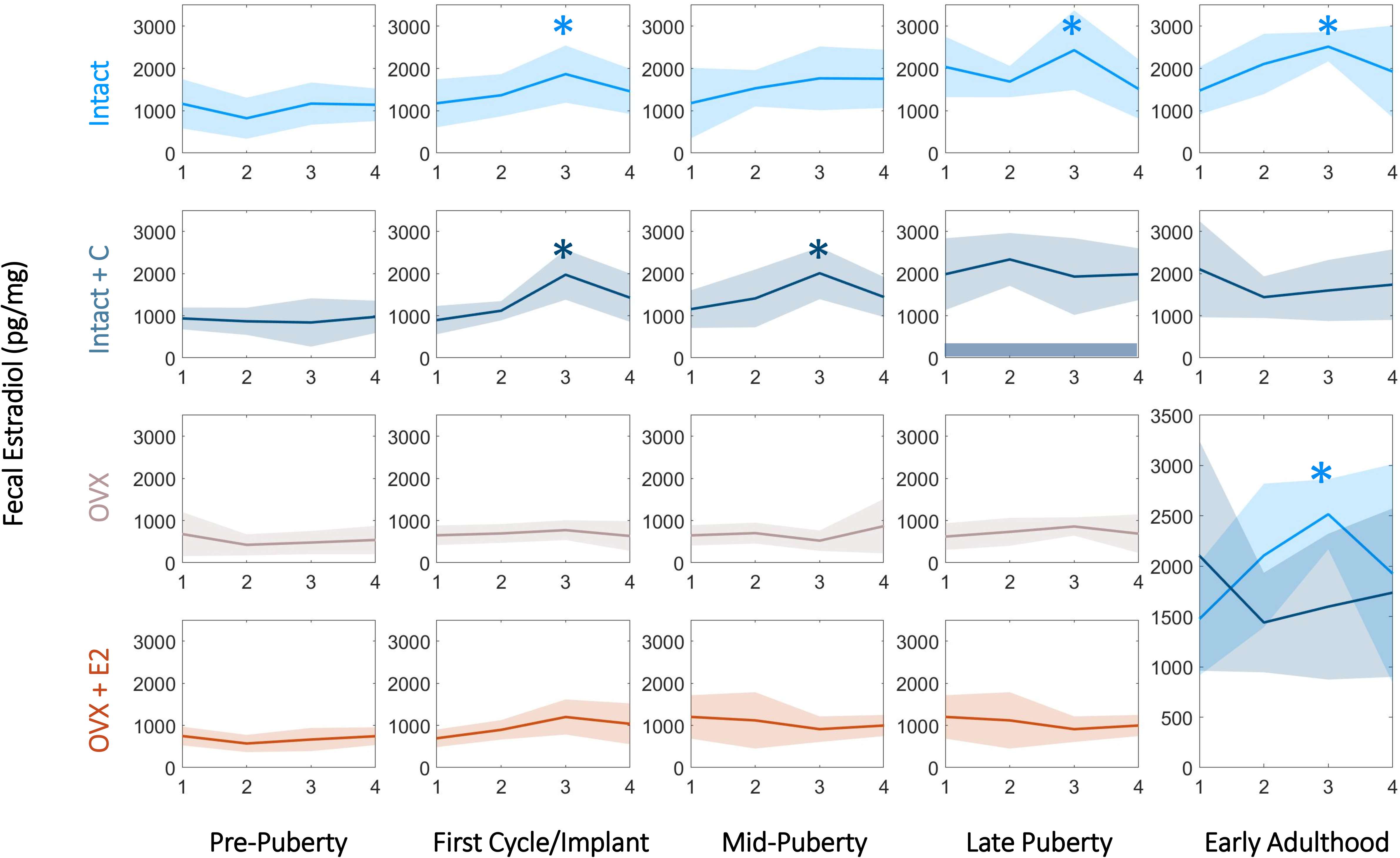
Estradiol Group Averages from Pre-Puberty to Adulthood Post-Contraception: Contraceptives Longitudinally Alter the Pattern but Not Level of Fecal Estradiol. Pre-puberty (p24-p30), early puberty (p30-p37), mid puberty (p43-p49), late puberty (p55-p61) and early adulthood (p69-p76) in Intact (A), Intact+C (B), OVX (C) and OVX+E2 (D). Early adulthood post-contraceptive administration, captured only in Intact and Intact+C animals, are overlaid in (E), illustrating persistent perturbation of estrous-cycle estradiol patterning. Dark bar indicates samples were collected during contraceptive administration in Intact+C animals. * Indicates significant elevation of day 3 compared to day 1 (p<0.05).

**Supplemental Figure 2.**
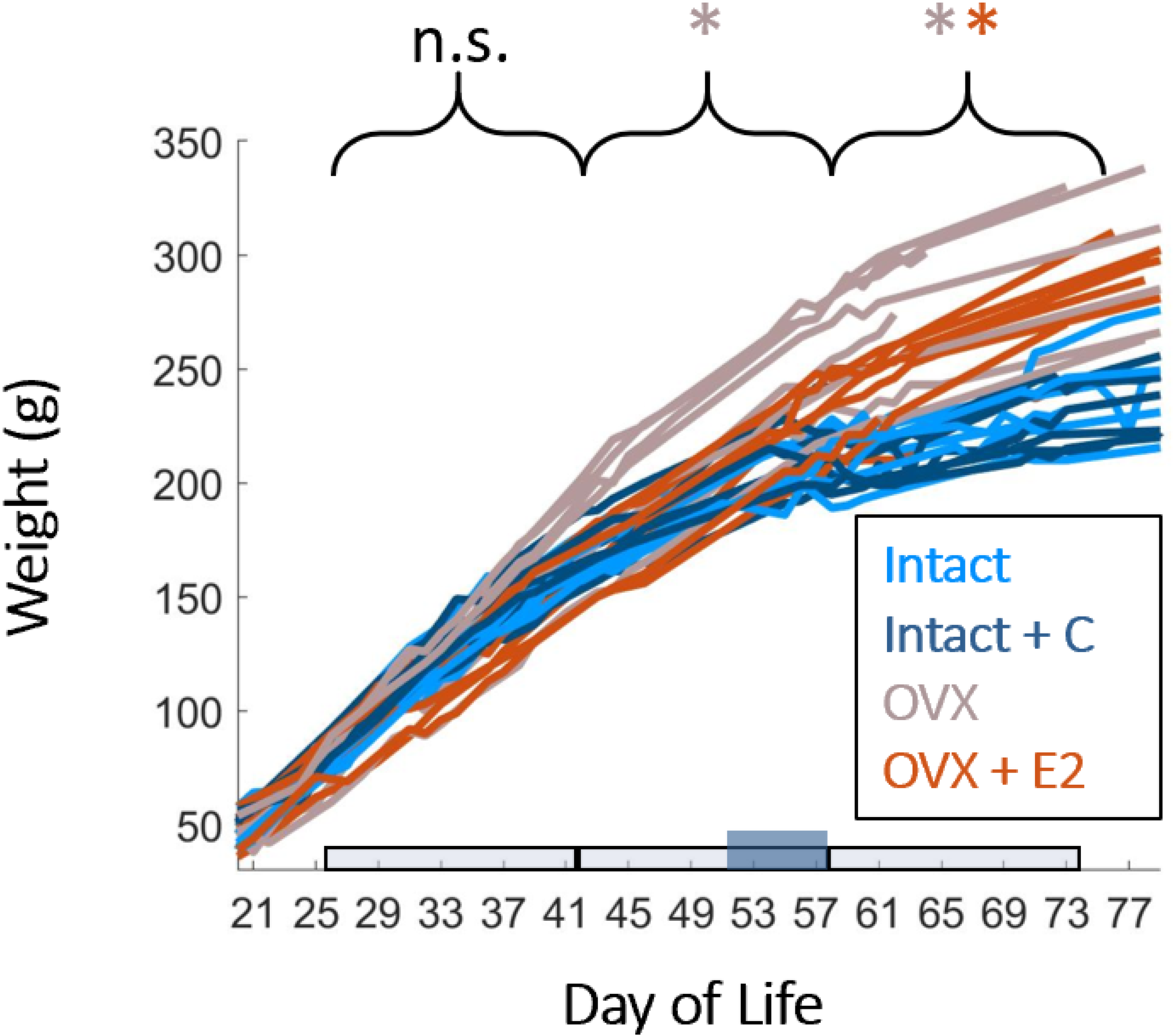
OVX and OVX + E2 are Heavier than Intact Animals in Mid Puberty and Early Adulthood. Prepubertal OVX did not significantly impact weight gain before mid-puberty. OVX animals (gray) are significantly heavier than all other groups at mid-puberty. OVX animals remain significantly heavier into adulthood. In early adulthood, OVX+E2 animals (orange) are not different from OVX animals and are significantly heavier than either Intact (light blue) or Intact+C animals (dark blue). Colors of * Indicate groups that are significantly heavier (p<0.05).

**Supplemental Figure 3.**
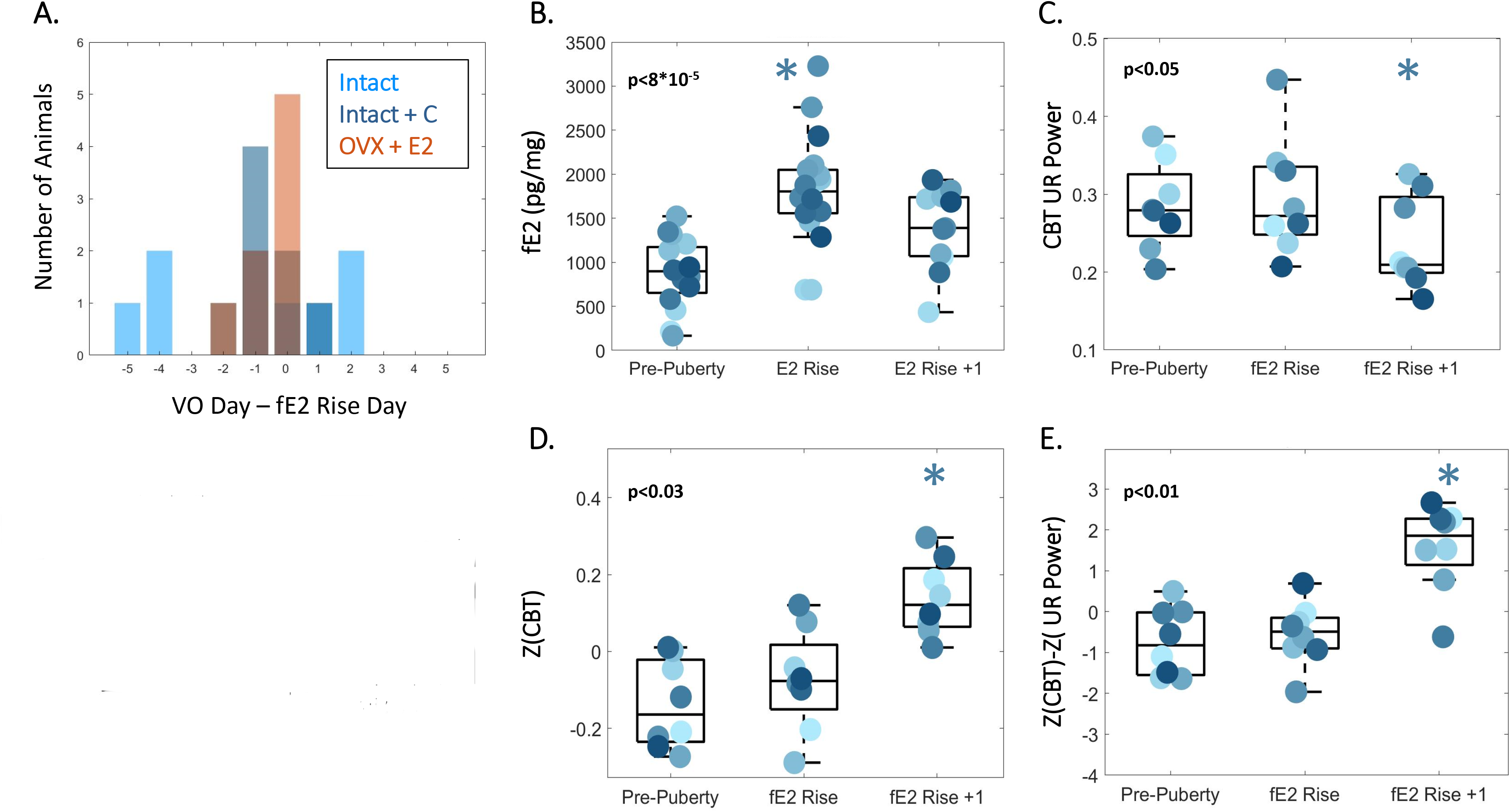
Day of First Estradiol Rise Occurs Near Vaginal Opening and Coincides with Markers of Estrus: Subsequent Drop in UR Power and Rise in CBT. Estradiol has a unique day in proximity to, but distinct from, vaginal opening, during which fE2 rises > 2 Standard Deviations within an individual (A-B). Alignment to this day as a proxy for puberty onset provides a convenient point from which to average across animals to permit easy visualization of the estrous cycle. Daily mean UR power decreases significantly following this day (C), and daily mean CBT rises significantly following this day (D), The difference between these two metrics exaggerates the difference, and the differences across subsequent cycles (E). After this point, all three metrics commence the 4-day estrous pattern. * Indicates significant difference from all other time periods.

**Supplemental Figure 4.**
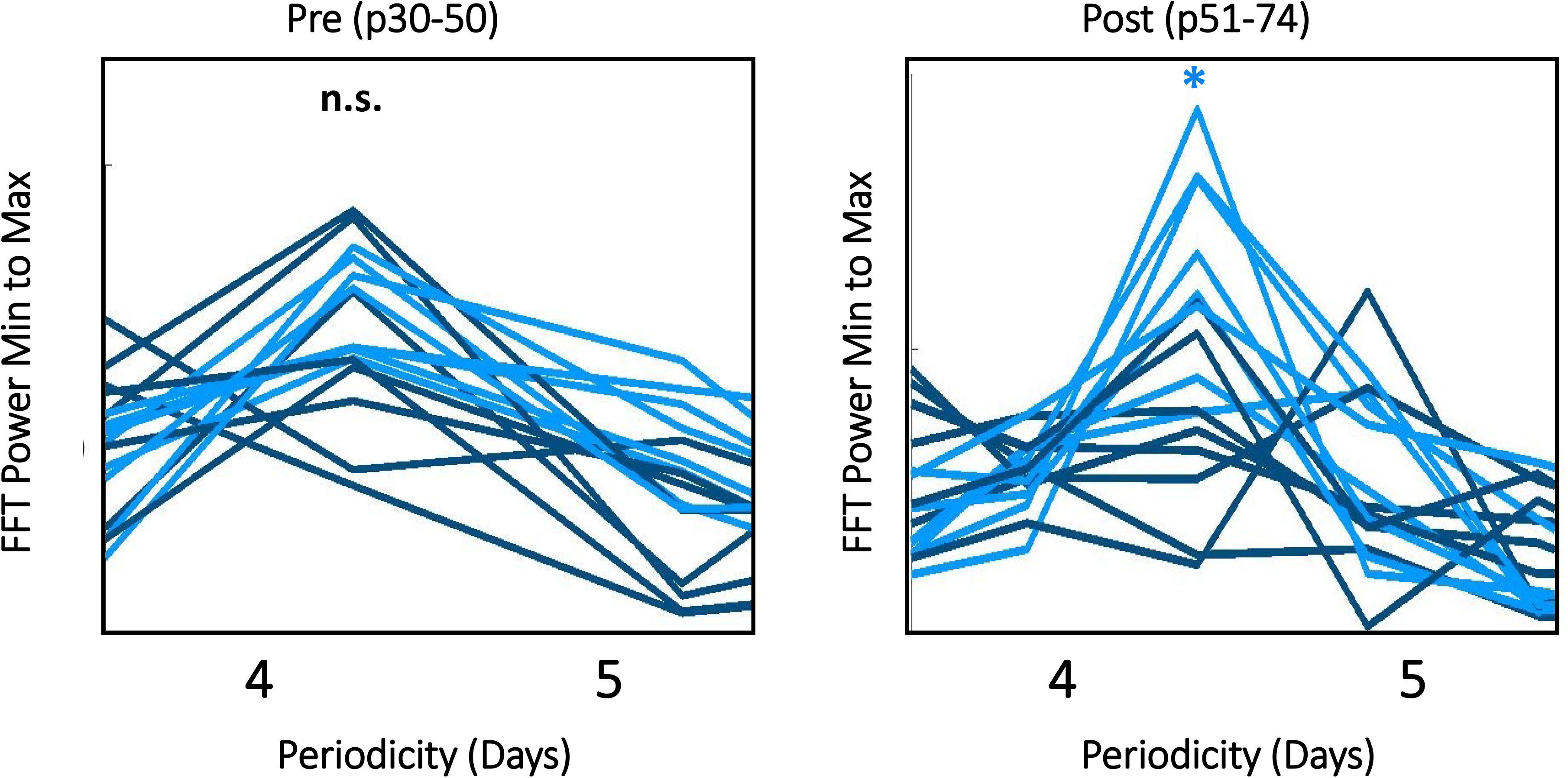
FFT Power of Intact Temperature Across Life Peaks Unique at 4 Days and is Disrupted During and After Contraceptive Administration. Fast Fourier Transform of Each Individual’s CBT Log in Intact (light blue) and Intact + BC (dark blue) prior to contraceptive administration (left), and during and after contraceptive administration (right). * Indicates significant difference for AUC between the groups in 4-5 periodicity.

**Supplemental Figure 5.**
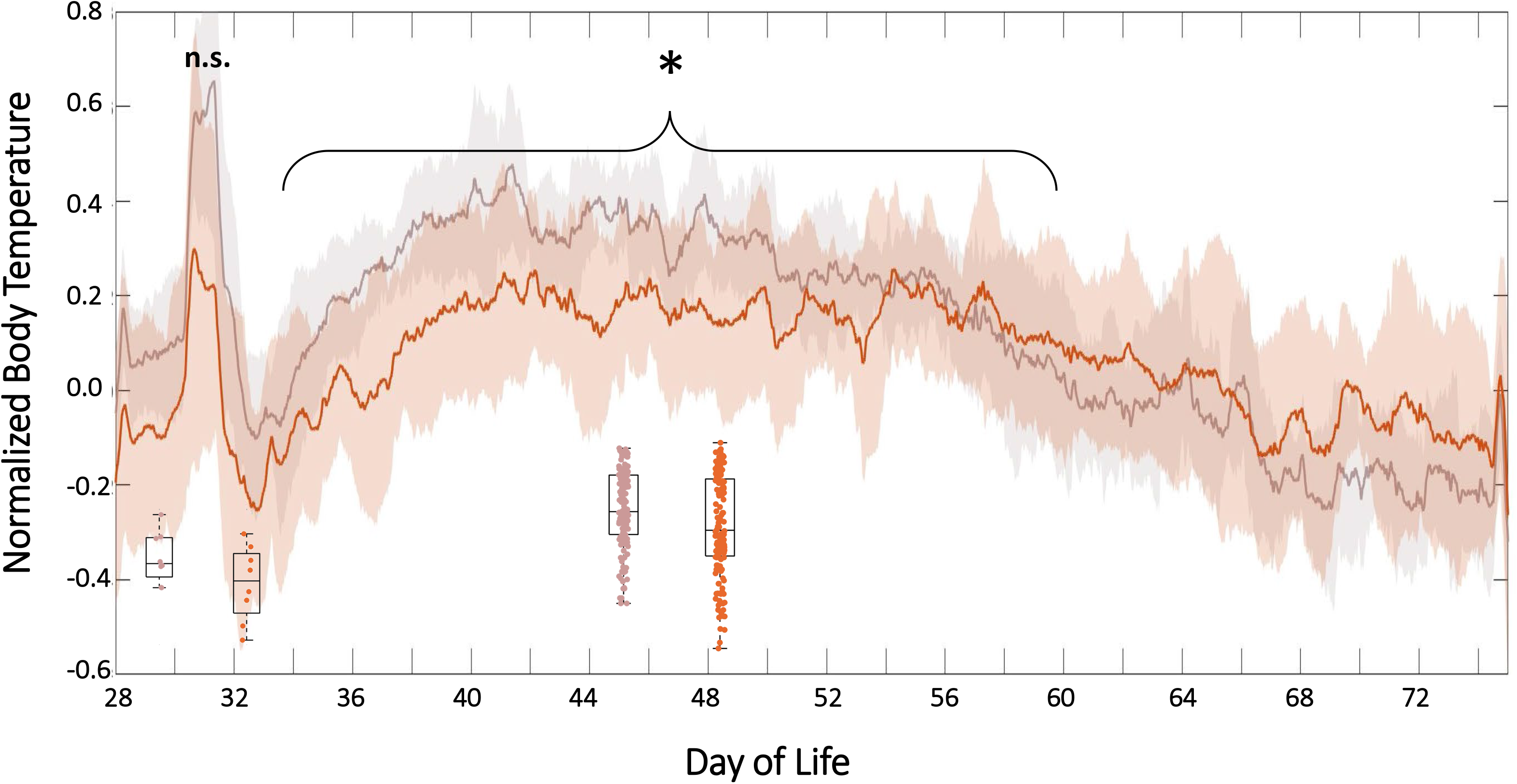
Estradiol Administration in Ovariectomized Animals is Associated with Lower Temperatures Across Adolescence. Core CBT linear group means (± standard deviation) in OVX (gray) and OVX+E2 animals from p28-p72. Box plots indicate values used in KW comparisons between groups during silastic implant recovery (left) and across adolescence (right) with independent y-axes. * Indicates significant difference between groups during the marked time period (p<0.03).

**Supplemental Figure 6:**
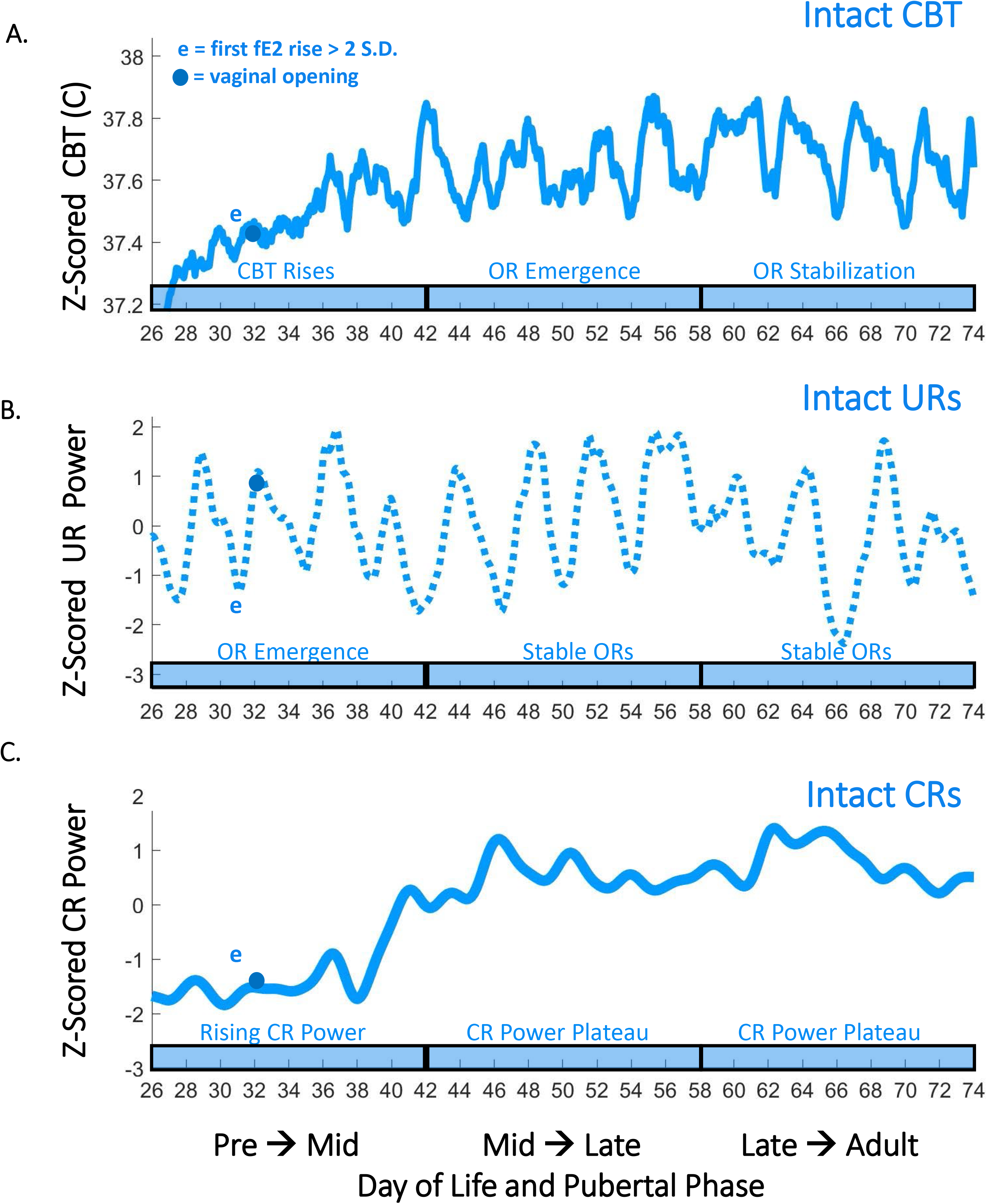
Summary of CBT Feature Use for Pubertal Staging in an Intact Female. Features of CBT and CBT rhythmicity can be used to complement and extend hormonal and external markers of adolescence. Here, we illustrate derived features on an example Intact female from p26 to p74, with fE2 rise and vaginal opening occurring on p31 and p32, respectively. Dots indicate vaginal opening and “e” indicates first rise of fE2 > 2 standard deviations. Rising CBT, emergence of ORs in UR power, and Rising CR power can be used to characterize pubertal onset. Mid to late adolescence is characterized by a plateau of CR power, as well as the emergence of ORs in raw CBT their persistence in UR power. Post-natal day 60 did not correspond with any overt transitions in CBT or CBT metrics.

## Notes

### Competing Interest Statement

The authors have declared no competing interest.

